# Constitutive NF-κB Activation is Amplified by VSV in Aggressive PC3 Prostate Cancer Cells that Resist Viral Oncolysis

**DOI:** 10.1101/2025.09.23.678061

**Authors:** Alaa A. Abdelmageed, Jack Smerczynski, Lute Douglas, Tori Russell, Matthew Morris, Stephen Dewhurst, Maureen C. Ferran

## Abstract

Cancer cells often have defects in antiviral pathways, making them susceptible to oncolytic viruses like vesicular stomatitis virus (VSV). However, some cancer cells resist viral infection through the constitutive expression of interferon-stimulated genes. This study examined whether NF-κB activation and NF-κB-dependent antiviral signaling contributes to resistance to VSV infection in the PC3 cell line, derived from an aggressive metastatic prostate cancer (PrCa) tumor. We found that NF-κB localized to the nucleus in VSV-infected PC3 cells, but not in the VSV-susceptible LNCaP PrCa cell line. Analysis of the upstream NF-κB inhibitor IκB-*α* revealed higher levels of both total and phosphorylated IκB-*α* in PC3 cells compared to LNCaP cells, indicating constitutive activation of the NF-κB pathway via an IκB-α–dependent mechanism. Notably, VSV infection did not alter IκB-α phosphorylation in PC3 cells, suggesting that VSV may amplify NF-κB signaling through an IκB-*α*–independent pathway. Furthermore, PC3 cells displayed elevated levels of the NF-κB p65 protein subunit compared to LNCaP cells, with its phosphorylated form significantly increased upon VSV infection. These results from phosphorylation assays confirm that multiple steps in the NF-κB pathway are differentially activated in PC3 and LNCaP cells. Additionally, the expression of several NF-κB-dependent cytokine and proinflammatory genes, including IL12 and IL6, were upregulated following VSV infection in PC3 cells, as compared to LNCaP cells. Blocking the NF-κB pathway using a pharmaceutical inhibitor resulted in increased PC3 cell death with VSV infection. Collectively, these findings suggest that enhanced NF-κB signaling may underlie the resistance of PC3 cells to VSV oncolysis, potentially offering new insights into therapeutic strategies targeting NF-κB in resistant prostate cancers.

## INTRODUCTION

Genetic abnormalities can accumulate in tumors, driving uncontrolled cell proliferation and survival. These mutations often perturb antiviral responses, including the interferon (IFN) pathway, leaving malignant cells susceptible to oncolytic (cancer-killing) viruses such as VSV (1-4). In contrast, healthy cells retain intact antiviral pathways that restrict viral replication and prevent cytotoxicity. Therefore, oncolytic viruses selectively replicate in cancer cells, causing direct cytolysis and the release of tumor-associated antigens, which trigger immunogenic cell death. This process enhances anti-tumor immunity by activating both innate and adaptive immune responses, promoting systemic tumor clearance and potential long-term therapeutic benefits (5).

Preclinical studies indicate that vesicular stomatitis virus (VSV) exhibits strong anti-tumor activity against various cancers, including lung (6-8), breast (9-11), and prostate cancers (PrCa) (12-14). Early-phase clinical trials have demonstrated that VSV-based oncolytic virotherapy is generally well-tolerated and holds promise as a therapeutic strategy, especially when combined with immune checkpoint inhibitors (15-17). Common approaches to improve safety and enhance tumor specificity of VSV include the use of recombinant viruses that are unable to suppress antiviral pathways in infected cells, a function of the viral M protein. These include viruses that express IFN type-I from the viral genome or contain a deletion or substitution at position 51 of M (18-20). In addition, efforts to increase targeting of the virus to cancer cells include combining VSV infection with different classes of chemotherapeutic drugs (20).

Despite these encouraging results, challenges remain. Resistance to oncolytic virus-mediated cytotoxicity has been observed across various cancer types and is considered a main challenge to developing viruses as oncolytic agents. Due to tumor heterogeneity, both virus-sensitive and virus-resistant cells can coexist in the same tumor (4). Resistant tumor cells exhibit reduced susceptibility to viral infection and replication; however, a comprehensive understanding of the molecular mechanisms that drive resistance is lacking.

Retention of antiviral pathways is believed to be a key factor in the resistance to oncolytic viruses. This has been observed in the PC3 PrCa cell line, which was derived from an aggressive metastatic PrCa tumor (21). PC3 cells exhibit reduced susceptibility to killing by both wild-type (WT) (13, 18, 22-24) and mutant VSV strains (13, 22, 25). Comparatively, the LNCaP cell line is sensitive to VSV infection (4, 13, 26) and was derived from a lymph node that was proximal to a prostate tumor (27). Another study demonstrated that multiple early stages of the VSV replication cycle are delayed in the PC3 cell line, but not in LNCaP cells, likely as a consequence of preserved antiviral signaling mechanisms (4).

Constitutive activation of NF-κB—a pivotal transcription factor involved in the induction of antiviral responses—has been observed in many aggressive and metastatic tumors (28). This constitutive activation is implicated in promoting several oncogenic processes, including accelerated cell cycle progression, increased cellular proliferation (29, 30), resistance to apoptosis, enhanced metastatic potential in prostate cancer (PrCa) (31), and reduced responsiveness to conventional therapies (32). Given these roles, the present study aimed to investigate whether NF-κB activation and NF-κB–dependent antiviral signaling contribute to the resistance of PC3 cells to oncolytic viral infection. The prognosis for patients with advanced metastatic PrCa remains poor, with a five-year survival rate of approximately 30% (13). Moreover, therapeutic options for late-stage disease are limited, and treatments effective in other solid tumors have shown minimal efficacy in this context. These challenges underscore an urgent need to develop novel therapeutic strategies for the management of advanced PrCa.

Our findings confirm that PC3 cells are resistant to VSV-mediated killing (13) and exhibit constitutive activation of NF-κB through an IκB-*α*–dependent pathway (28, 33). In contrast, LNCaP cells are susceptible to VSV infection (4, 13, 26) and do not exhibit constitutive NF-κB activity (26). In PC3 cells, phosphorylation of IκB-α at serine residue 32 was unaffected by VSV infection, suggesting that the constitutive activity of NF-κB is not further modulated through canonical IκB-*α* phosphorylation. Notably, PC3 cells displayed higher levels of the NF-κB p65 protein subunit relative to LNCaP cells, with the most elevated levels of phosphorylated p65 (serine 276) following VSV infection.

NF-κB translocation in PC3 cells was found to be significantly higher than in LNCaP cells in the absence of viral infection, indicating constitutive activation of NF-κB in the resistant cells. This was associated with high baseline levels of both total and phosphorylated IκB-α. Following VSV infection in PC3 cells, nuclear translocation (activation) of NF-κB was further increased, but IκB-α phosphorylation was unchanged, suggesting that VSV may amplify NF-κB signaling in PC3 cells through an IκB-*α*–independent/non-canonical pathway. In contrast, little NF-κB activation was noted in LNCaP cells, regardless of VSV infection. To evaluate downstream functional consequences of this differential NF-κB signaling, we assessed the expression of NF-κB–dependent effector genes in response to VSV infection. Baseline expression of several genes encoding antiviral cytokines and proinflammatory mediators was higher in uninfected PC3 cells compared to LNCaP cells, and VSV infection further amplified their expression.

Collectively, these findings suggest that constitutive and VSV-enhanced NF-κB activation contributes to the resistance of the highly metastatic PC3 PrCa cell line to VSV infection, likely through the upregulation of NF-κB–dependent antiviral and inflammatory signaling pathways.

## METHODS

### Cells, viruses, and infections

The LNCaP and PC3 human PrCa cell lines were obtained from the American Type Culture Collection (ATCC) (CRL-1740 and CRL-1435, respectively) and were grown in aRPMI-1640 containing 10% FBS (ATCC). The heat-resistant (HR-C) strain of the Indiana serotype of VSV was used as the WT virus. Mutant T1026R1 (R1), a temperature-stable revertant of T1026, was derived from the Indiana HR-C strain of VSV by chemical mutagenesis (supplied by C. P. Stanners). All viruses were grown on Vero cells as described (34) in EMEM growth medium (ATCC). Cells were infected with each virus at multiplicities of infection (MOI) ranging from 0.1, 1, 5, 10, to 50 plaque forming units (PFU)/cell (see Figure legends). Viruses were allowed to adsorb to target cells in serum-free culture medium for 1 hour at 37 °C, after which complete (serum-containing) culture medium was added to the cells.

### PrestoBlue Viability Assay

PC3 and LNCaP cells were seeded in Santa Cruz Biotech black frame/clear bottom cell culture 96-well plates (sc-204468). After 24 hours, cells were infected with either WT VSV or the R1 mutant at an MOI of 0.1 or 1. At 24, 48 and 72 hours post infection (hpi), supernatants were removed and 100 µL of master mix was added, composed of 10% PrestoBlue HS Cell Viability Reagent (Thermo Fisher Sci) in serum-free, phenol red-free RPMI medium. This viability reagent is reduced by viable cells and converted into a red fluorescent dye. Plates were then incubated in the dark at 37°C for 40 minutes and a fluorescence reading was taken on a SpectraMax iD3 Plate Reader (Molecular Devices) using an Em/Ex 590/550 nm wavelength range.

### Immunofluorescence

Immunofluorescence analysis was conducted as described (34, 35) except that cells were blocked with UltraCruz® Blocking Reagent (sc-516214, Santa Cruz Biotechnology) for 30 minutes followed by incubation with an NF-κB-(p65) antibody conjugated to Alexa Fluor® 488 (sc-8008 AF488, Santa Cruz Biotechnology) for 90 minutes at room temperature. Nuclear localization of p65 was quantified using ImageJ. Locations of cell nuclei and density of NF-κB staining in those regions were determined by automated analysis of DAPI and FITC channel images. Each DAPI channel image was subjected to a Gaussian blur filter with a sigma value of 2. Next, the image was thresholded using the default algorithm, and the watershed tool was used to separate adjoining nuclei. Finally, nuclei were defined as regions of interest ranging from 200-20,000 pixels in size, with nuclei on the edge of each image excluded. The intensity of NF-κB staining in each nuclear region was then quantified. The background fluorescence times the area of each nuclear region was subtracted from the integrated fluorescent density of NF-κB staining in each nucleus to yield the corrected total cell fluorescence (CTCF) for each cell. Median CTCF among all cells in each image was taken as representative.

### IκBα LUMIT Assay

LNCaP or PC3 cells in Corning white 96-well cell culture plates were treated with 20 µM (30 µl) of the proteasome inhibitor MG132 for 1 hour before infection. Next, PC3 cells were infected with WT or R1 VSV at an MOI of 50 PFU/cell for 1 hour, while LNCaP cells were infected at an MOI of 10 PFU/cell. After adsorption, the cells were re-fed with complete media containing 20 µM (40 µl) of MG132. At the indicated hpi, the amount of total and phosphorylated IκBα was determined using the LUMIT Immunoassay Cellular Systems Kit from Promega according to the manufacturer’s directions. Briefly, the cells were lysed, and the appropriate combination of primary and secondary antibodies were added to the cell lysates. Luminescence substrate was introduced to the antibody lysate mixture and luminescence values of phosphorylated and total IκBα protein were quantified using a SpectraMax iD3 Plate Reader (Molecular Devices).

### P65 LUMIT Assay

LNCaP or PC3 cells were seeded in a Corning white 96-well cell culture plate 24 hours prior to infection. Cells were infected with WT or R1 VSV at an MOI of 50 PFU/cell for PC3 cells and an MOI of 10 PFU/cell for LNCaP cells. At the indicated hpi, the amount of total and phosphorylated p65 protein was determined using the LUMIT Immunoassay Cellular Systems Kit from Promega according to the manufacturer’s directions. Briefly, the cells were lysed, and the appropriate combination of primary and secondary antibodies were added to the cell lysates. Luminescence substrate was introduced to the antibody lysate mixture and luminescence values of phosphorylated and total p65 protein were quantified using a plate reader.

### RNA Sequencing

Cells were grown in 30 mm tissue culture dishes coated with poly-D-lysine (PDL). Infection was carried out using VSV at an MOI of 50 and 10 PFU/cell for PC3 and LNCaP, respectively. At 6 hpi for LNCaP and 8 hpi for PC3 cells, cells were lysed in plates using RIPA lysis buffer (Thermo Scientific RIPA Lysis and Extraction Buffer). Samples were sent to the University of Rochester Genomic Center for library preparation and sequencing. Analysis was performed on count data using the Bioconductor DESeq2 package to calculate differential expression (DE) levels. Contrasts were made on the DE data by comparing virus treatments to mock treatment for each cell line.

### Statistical Analysis

Statistical analysis throughout this paper was performed in R using either the student’s t test or the one-way ANOVA test and an asterisk indicates a significant change (p<0.05). Unless otherwise noted, results are expressed as means and error bars indicate the ± standard error of the means (SEM).

## RESULTS

### PC3 cells are more resistant to VSV-mediated killing than LNCaP cells

To assess cellular resistance to VSV, cell death in response to infection with WT VSV or the mutant VSV strain, R1, at an MOI of either 0.1 or 1 MOI was determined using a PrestoBlue assay. PC3 cells exhibited lower levels of cell killing, as compared to LNCaP cells at 24, 48, and 72 hpi (Fig. 1). This was especially evident at 72 hpi with WT VSV at an MOI of 0.1, and at 72 hpi with R1 VSV at an MOI of 1. These findings confirm the greater resistance of PC3 cells to VSV-mediated cell killing, versus LNCaP cells.

**Figure 1:**
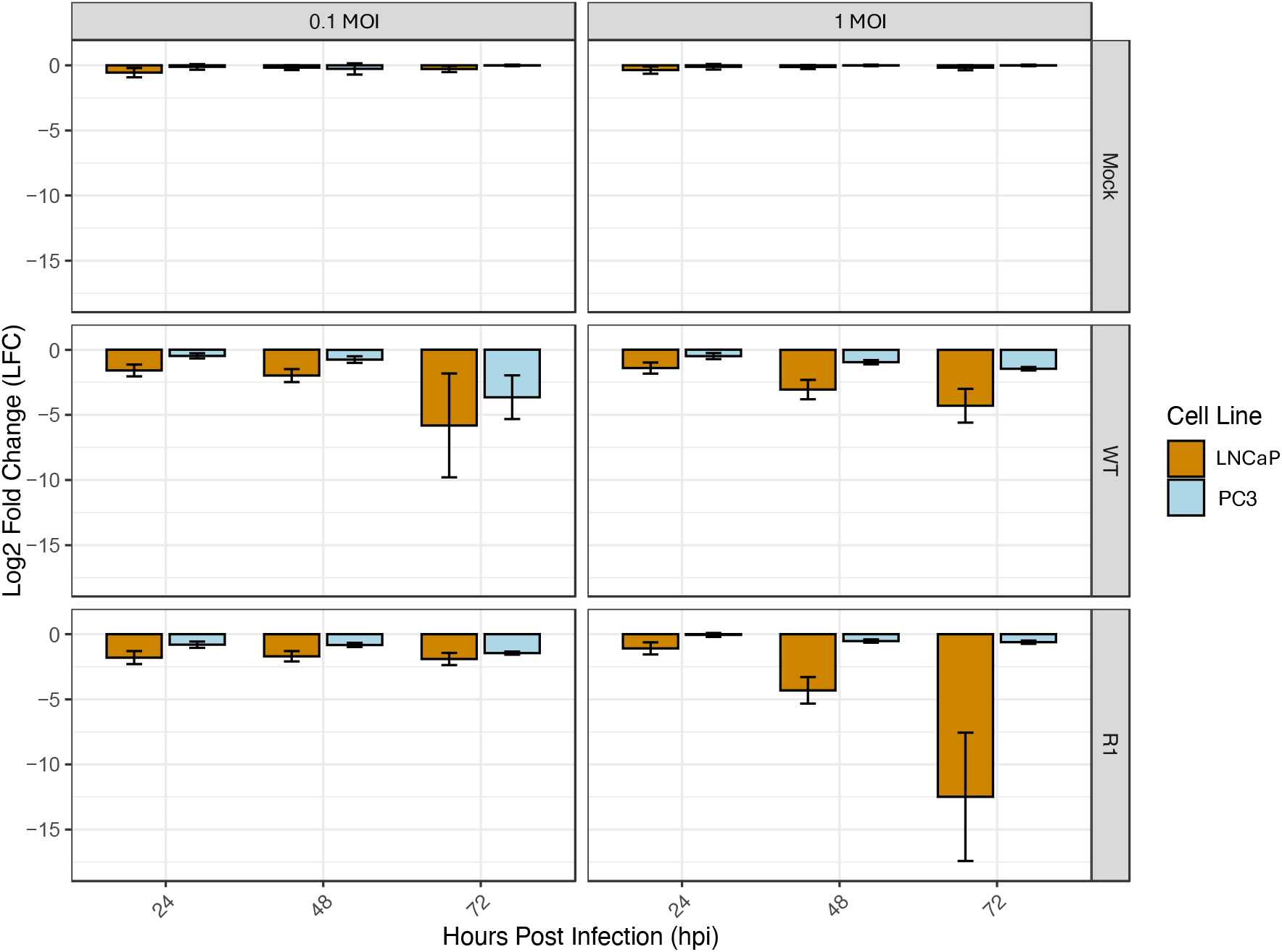
PC3 cells are more resistant to VSV-mediated lysis than LNCaP cells. PC3 and LNCaP cells were infected with two VSV strains: WT and the R1 mutant strain, at an MOI of 0.1 and 1. Cell death was then measured at 24, 48 and 72 hours using a PrestoBlue assay. Statistical analysis was performed on three rounds of experiments, each with three biological replicates for each treatment/timepoint. Log2 Fold Change (LFC) was calculated relative to mock treated cells at each of the timepoints and treatments.

### Baseline NF-κB nuclear translocation is elevated in PC3 cells relative to LNCaP cells, and is further increased by VSV infection

To test whether VSV infection of the two cell lines leads to activation of the proinflammatory and antiviral NF-κB pathway, an immunofluorescence (IF) assay of NF-κB nuclear localization was performed. As a positive control, cells were treated with TNF-*α*, a potent activator of NF-κB which resulted in NF-κB nuclear localization in virtually all PC3 cells, but in relatively few LNCaP cells (Fig. 2A) – suggesting that both cell types are capable of responding to signals that induce NF-κB activation, but that PC3 cells do so more strongly than LNCaP cells. Consistent with this, nuclear localization of the NF-κB subunit, p65, was significantly higher in mock-treated PC3 cells than in mock-treated LNCaP cells (Fig. 2B), indicating elevated baseline levels of NF-κB activation in PC3 cells versus LNCaP cells.

To address the effect of VSV infection on NF-κB nuclear localization, cells were infected with either WT VSV or the R1 VSV mutant. For these experiments, a lower MOI of virus was used to infect the highly susceptible LNCaP cells (MOI 10) versus the more resistant PC3 cells (MOI 50), in order to achieve synchronous viral infection (13) while avoiding excessive cell death over the course of the experiment. The levels of NF-κB nuclear localization were significantly elevated in PC3 cells following infection with both WT VSV and the R1 VSV mutant, as compared to uninfected cells (mock; Fig. 2B). In contrast, NF-κB nuclear localization in LNCaP cells was unaffected (or even modestly reduced) by VSV infection (Fig. 2B).

**Figure 2:**
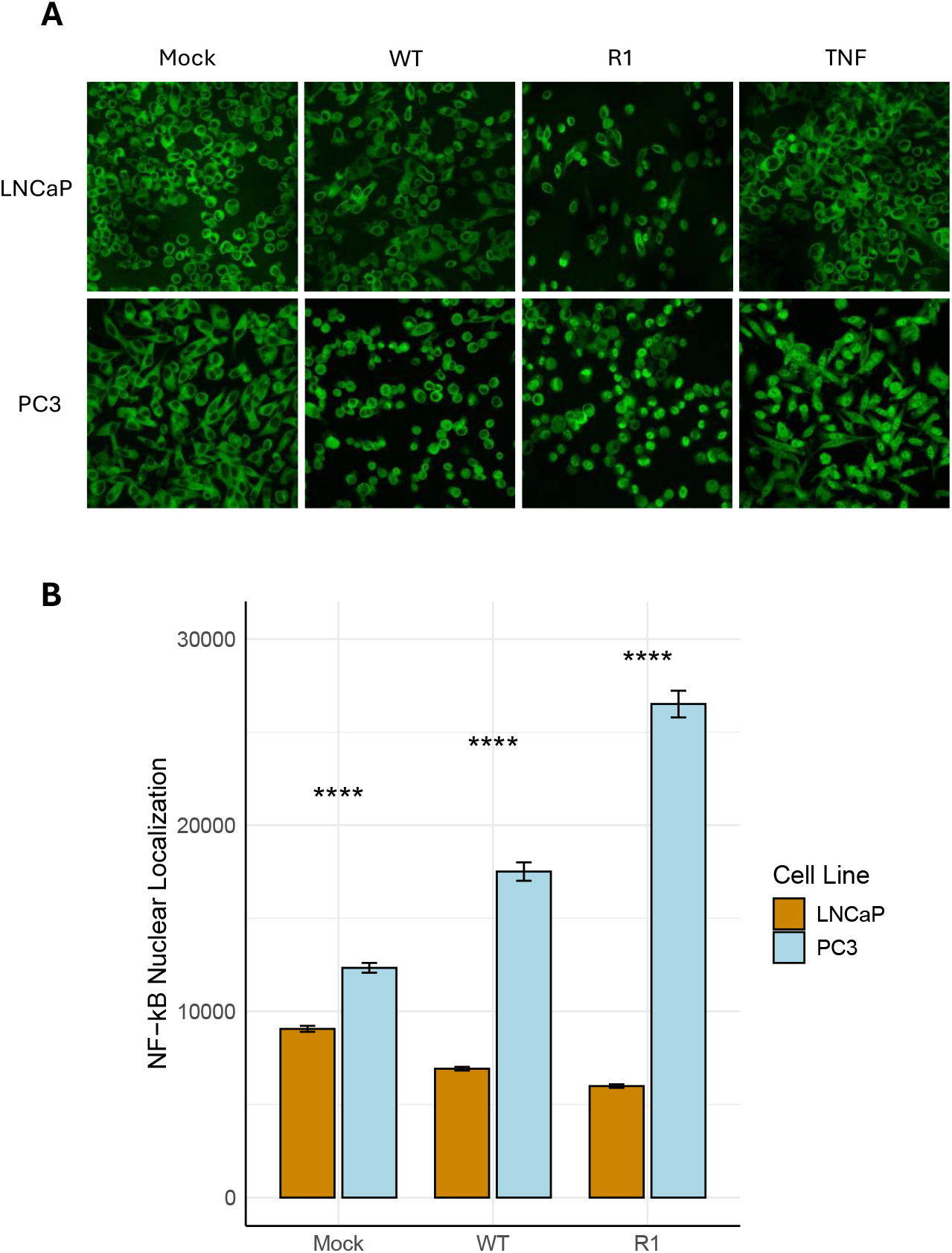
PC3 cells exhibit greater baseline and VSV-induced levels of NF-κB nuclear localization than LNCaP cells. A) Immunofluorescence images of NF-κB localization in PC3 and LNCaP cells, following treatment with TNF-*α*, mock treatment or infection with VSV (WT and R1) at an MOI of 50 (PC3 cells) or 10 (LNCaP cells). B) NF-κB nuclear localization was quantitated by automatic counting of corrected total cell fluorescence (CTCF) in ImageJ. Statistical analysis was performed on three rounds of experiments, each performed using five biological replicates for each treatment. 10 images for each biological replicate were loaded into the software for counting. Statistical analysis was performed in R using a paired student’s t-test to compare PC3 to LNCaP cells in each treatment category (p < 0.001: ***; p < 0.01: **; p < 0.05: *).

### Total and phosphorylated levels of the NF-κB-inhibitor, IκB-*α*, are higher in PC3 cells than in LNCaP cells, under all conditions tested (including baseline)

To understand the underlying basis for the higher levels of NF-κB nuclear translocation in PC3 cells, we examined the levels of expression and phosphorylation of IκB-*α*, an inhibitory protein that binds to NF-κB and retains it in the cytoplasm. The levels of both total and phosphorylated IκB-*α* were measured by LUMIT assays. In all conditions (including baseline), and at all time points examined, total levels of IκB-α were higher in PC3 cells than in LNCaP cells, (Figure 3). Phosphorylated levels of IκB-α were also significantly higher in PC3 cells under all conditions (including baseline). This is important because phosphorylation of IκB-α is associated with accelerated degradation of this inhibitory protein - and thus higher levels of NF-κB activation.

### Total levels of the NF-κB subunit, p65, are higher at baseline in PC3 cells than in LNCaP cells

The contribution of the NF-κB subunit p65, or RelA, to the differential activation of the NF-κB pathway in PC3 versus LNCaP cells was determined by using LUMIT assays to measure total and phosphorylated levels of p65 in both cell types, both at baseline and in response to VSV infection. This assay focused on phosphorylation of serine 276 of p65, since this is known to induce a conformational change that increases its interaction with transcription coactivators like CREB binding protein (CBP) - thereby enhancing the transcriptional activity of NF-κB (36). At baseline, total levels of p65 were higher in PC3 cells than in LNCaP cells (Fig. 4). Moreover, levels of phosphorylated p65 were elevated in VSV-infected PC3 cells compared to VSV-infected LNCaP cells (especially in response to infection with WT VSV (Fig. 4)). Therefore, the constitutive NF-κB activation observed in PC3 cells (Fig. 2) is likely due to increased IκBα phosphorylation (Fig. 3) and elevated total expression of the p65 subunit of NF-κB (Fig. 4).

**Figure 3:**
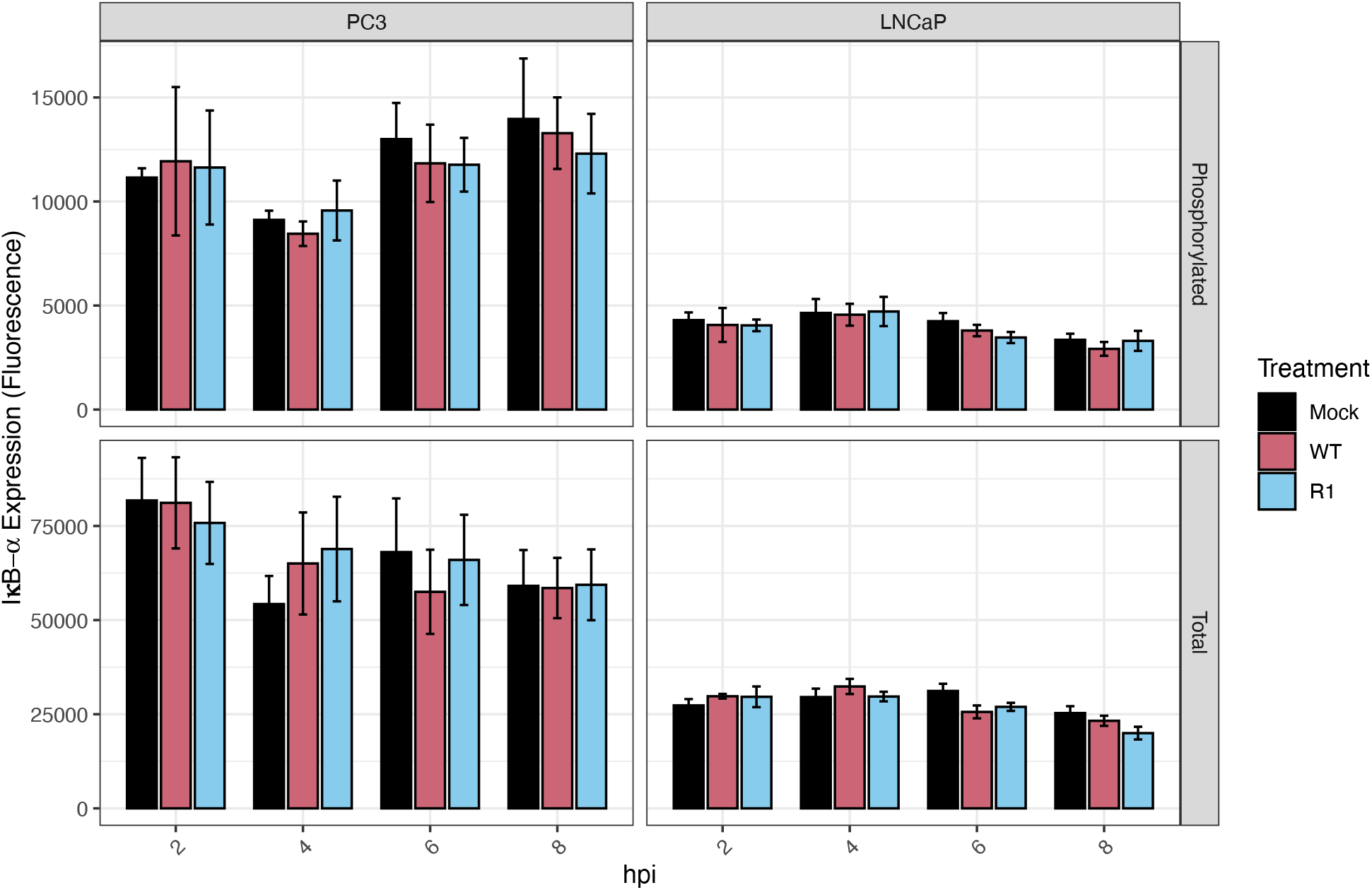
Total and phosphorylated levels of the NF-κB-inhibitor, IκB-*α*, are higher in PC3 cells than in LNCaP cells, under all conditions tested. A LUMIT assay was performed to measure total and phosphorylated levels of IκB-*α* in PC3 and LNCaP cells (normalized to cell count), following mock infection or infection with VSV (WT and R1) at an MOI of 50 (PC3 cells) or 10 (LNCaP cells); samples were analyzed at the indicated time points (hours post-infection, hpi). Statistical analysis was performed on three rounds of experiments, with three biological replicates for each treatment. The Figure was generated in R and error bars were integrated by calculating the standard error of means.

**Figure 4:**
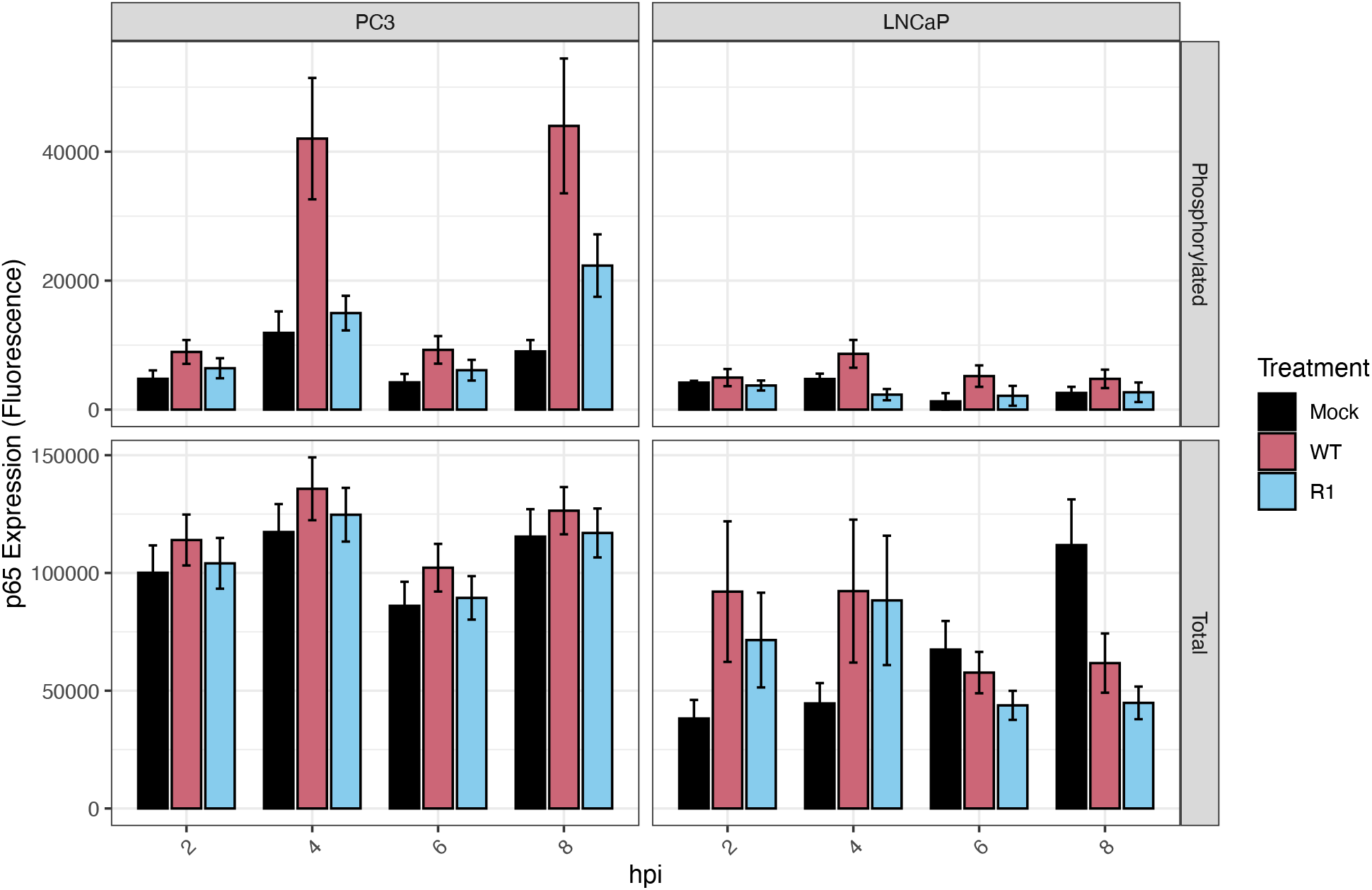
Total levels of the NF-κB subunit, p65, are higher at baseline in PC3 cells than in LNCaP cells. A LUMIT assay was performed to measure total and phosphorylated levels of p65 in PC3 and LNCaP cells (normalized to cell count), following mock infection or infection with VSV (WT and R1) at an MOI of 50 (PC3 cells) or 10 (LNCaP cells); samples were analyzed at the indicated time points (hours post-infection, hpi). Statistical analysis was performed on three rounds of experiments, with three biological replicates for each treatment. The Figure was generated in R and error bars were integrated by calculating the standard error of means.

### PC3 cells express higher levels of representative antiviral genes in response to VSV infection than LNCaP cells

The data in Figs. 2 and 3, indicate that baseline levels of NF-κB activation are elevated in PC3 cells versus LNCaP cells, and that PC3 cells express higher levels of phosphorylated p65 in response to VSV infection (Fig. 4). This raises the question of the levels of expression of NF-κB-dependent antiviral genes in the two cell types. Fig 5 shows representative expression data for selected antiviral/proinflammatory genes including both cytokines (IL6, IL12, IL15, TNF-*α*, IFN-*β*) and the chemokine CXCL8. VSV infection resulted in much higher levels of expression of all of the tested cytokine genes in PC3 cells, as compared to LNCaP cells (Fig. 5). In general, the R1 mutant of VSV was a more powerful activator of the expression of these genes than WT VSV (e.g., see data for CXCL8 and TNF-*α*). These results indicate that activation of the NF-κB pathway results in increased expression of proinflammatory and antiviral genes, including both cytokines (IL6, IL12, IL15, TNF-*α*, IFN-*β*) and the chemokine CXCL8, following VSV infection of PC3 cells, as compared to VSV-sensitive LNCaP cells (Fig. 5).

**Figure 5:**
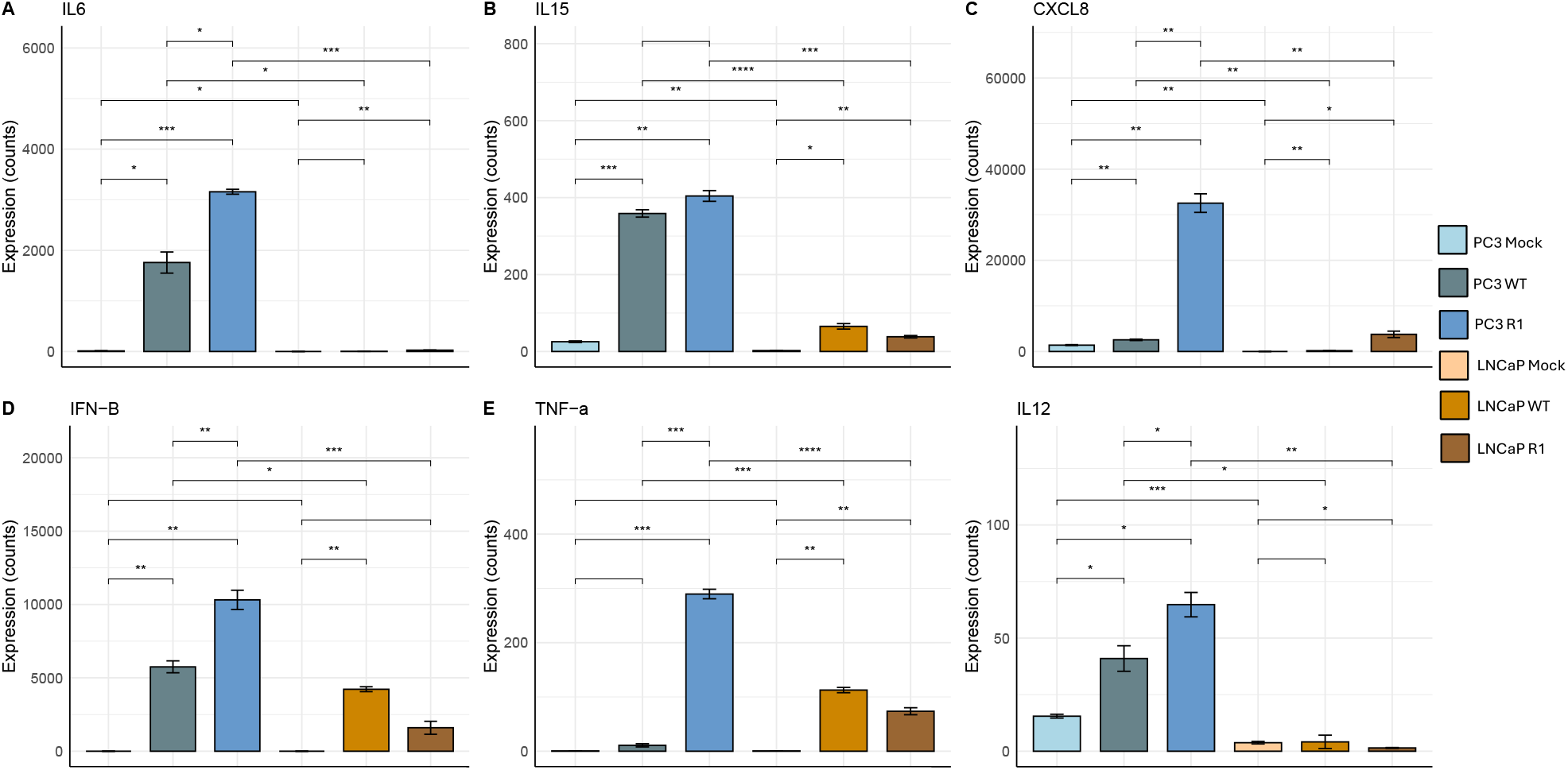
PC3 cells express higher levels of representative antiviral genes in response to VSV infection than LNCaP cells. mRNA expression levels for selected antiviral/proinflammatory genes were measured in PC3 and LNCaP cells, following mock infection or infection with VSV (WT and R1) at an MOI of 50 (PC3 cells) or 10 (LNCaP cells); samples were analyzed at 6 and 8 hpi for PC3 and LNCaP, respectively. Sequencing analysis was performed on three biological replicates for each treatment. Statistical analysis was performed in R using a paired student’s t-test comparing PC3 to LNCaP cells in each treatment category and error bars were integrated by calculating the standard error of means (p.value < 0.001: “***”, p.value < 0.01: “**”, p.value < 0.05: “*”).

### Cell survival steeply decreases in PC3 cells treated with Bay11-7082 upon VSV infection

To follow-up on our observations, we aimed to test whether blocking the NF-κB pathway would result in increased cell death of PC3 cells in the presence of VSV infection. Cells were treated with the NF-κB pathway blocker, Bay11-7082, at 2.5 uM for 24, 48, and 72 hours. Viral infection was carried out at 8 hours prior to each timepoint. Cell survival was unaffected by viral infection nor Bay11-7082 treatment alone (Fig. 6). However, samples both treated with Bay11-7082 and infected with VSV displayed a significant increase in cell death across all timepoints tested, indicating synergy. These results suggest that blocking the NF-κB pathway in PC3 cells renders them susceptible to killing by VSV.

**Figure 6:**
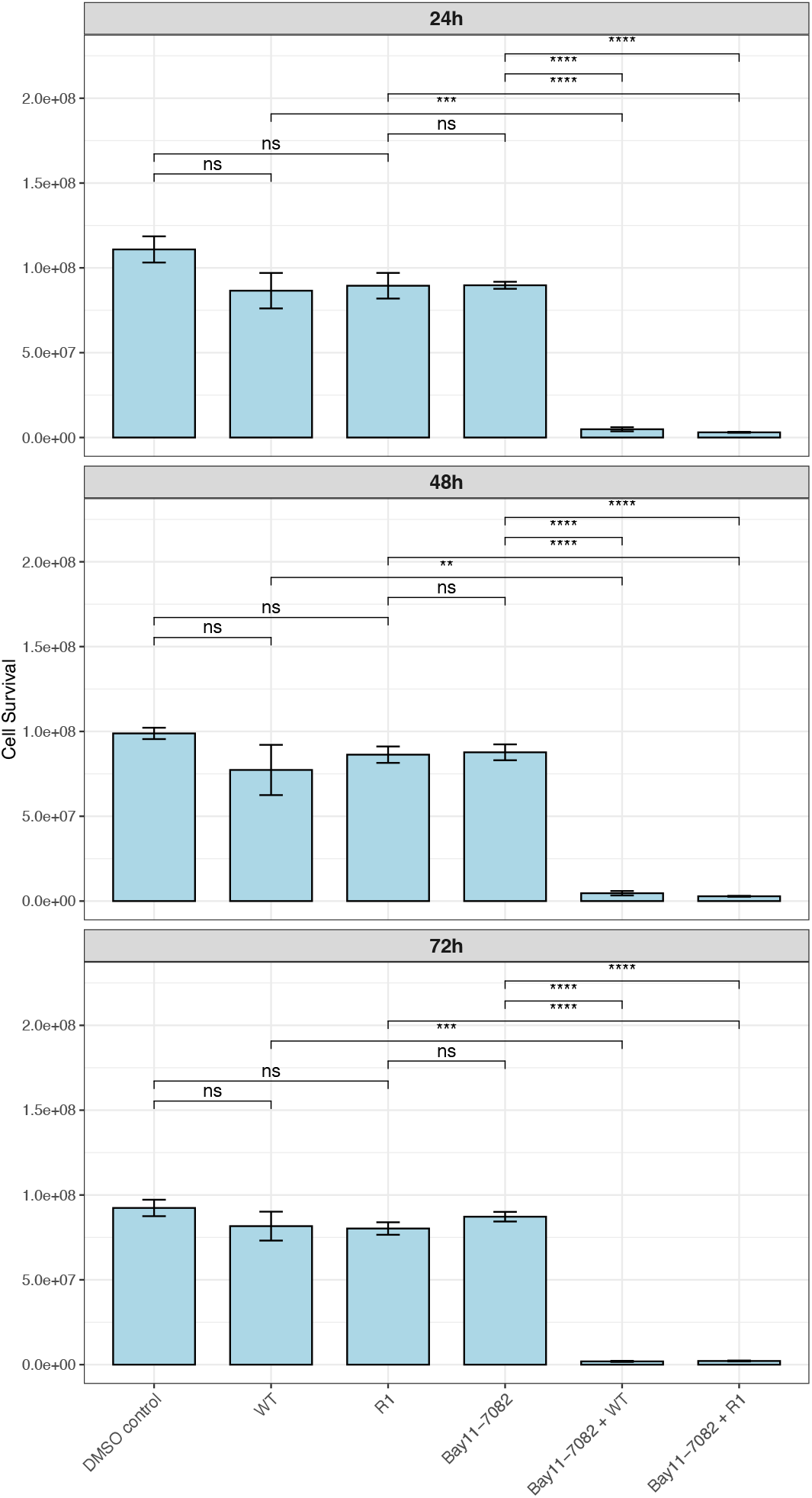
Blocking of NF-κB pathway results in cell death upon VSV infection in PC3 cells. Cell survival levels, with and without the NF-κB pathway inhibitor Bay11-7082, were measured in PC3 cells, following DMSO control treatment or infection with VSV (WT and R1) at an MOI of 50. Samples were run in duplicates and analyzed with PrestoBlue assay at 24, 48, and 72h post-treatment with Bay11-7082. Viral infection was initiated 8 hours prior to analysis at each timepoint. Statistical analysis was performed in R using student’s t-test comparing each treatment category and error bars were integrated by calculating the standard error of means (p.value < 0.0001: “****”, p.value < 0.001: “***”, p.value < 0.01: “**”, p.value < 0.05: “*”).

## DISCUSSION

The use of VSV in cancer therapeutics is challenged by resistance of certain cancer cell populations to oncolytic killing by VSV. Our findings implicate NF-κB signaling as a key determinant of resistance to VSV-mediated oncolysis in aggressive prostate cancer. Specifically, the constitutive and VSV-induced activation of NF-κB in PC3 cells supports an antiviral state that restricts VSV replication and cytolysis, whereas LNCaP cells, which lack baseline NF-κB activation, remain permissive to viral replication and are efficiently killed by VSV.

The implications of our findings may extend beyond prostate cancer, as constitutive NF-κB activation is a hallmark of numerous malignancies, including pancreatic (37), colorectal (38), breast (39), and certain hematologic cancers (40). In many types of tumors, persistent NF-κB signaling may contribute to therapy resistance by promoting cell survival, sustaining antiviral defenses, and shaping an immunosuppressive microenvironment (41). These mechanisms pose significant barriers to the efficacy of oncolytic virotherapy. Our results support the broader concept that NF-κB activity is a critical determinant of tumor susceptibility to oncolytic viruses and suggest that NF-κB inhibition could enhance virotherapy across multiple cancer types. Furthermore, these insights are likely applicable across a variety of other oncolytic viral platforms—including herpes simplex virus, reovirus, and adenovirus—which are similarly restricted by antiviral responses (42).

Given the observations presented here, therapeutic targeting of NF-κB represents a potential strategy to sensitize resistant prostate tumors to oncolytic virotherapy. However, the clinical development of NF-κB inhibitors is complicated by the pathway’s dual roles in cancer biology. Canonical NF-κB signaling, primarily mediated by the p65/p50 heterodimer, is frequently associated with tumor-promoting functions, including enhanced survival, proliferation, inflammation, and metastasis. NF-κB also transcriptionally regulates multiple anti-apoptotic genes such as BCL-2, BCL-xL, and cIAPs, which can contribute to therapy resistance (43). In this context, NF-κB inhibition could lower the apoptotic threshold of tumor cells, potentially enhancing the cytolytic activity of VSV and other therapeutic agents.

However, NF-κB’s role is not uniformly pro-oncogenic. In certain cellular and inflammatory contexts, NF-κB signaling can contribute to apoptosis or senescence, (44). Furthermore, the NF-κB family comprises five subunits (p65/RelA, c-Rel, RelB, p50, and p52) that participate in both canonical and non-canonical pathways, each with distinct target gene repertoires and biological outcomes. While the canonical p65/p50 pathway is often the primary driver of antiviral gene expression, contributions from alternative subunits—such as RelB/p52 in non-canonical signaling—should not be excluded, especially given their roles in tumor progression and immune evasion in other malignancies (45).

Our findings highlight the importance of a nuanced approach to NF-κB inhibition, particularly when employing approaches such as small molecule inhibitors of IKKβ, or treatment with pharmacological inhibitors of NF-κB (e.g. IKK16), that broadly suppress systemic NF-κB signaling, which is critical for innate immunity, inflammation, and tissue homeostasis (46, 47). Therapies that incorporate NF-κB inhibitors in a tumor-targeted fashion could synergize to overcome resistance in tumor cells, while avoiding systemic inhibition of NF-κB. One approach involves arming oncolytic viruses, such as recombinant VSV, with an NF-κB inhibitory gene (e.g., IκB*α* super-repressor, A20). This would limit expression of the inhibitor to infected tumor cells, thereby sensitizing resistant tumors without affecting NF-κB in surrounding normal tissue. Additionally, gene-silencing technologies such as CRISPR/Cas9 and RNAi can selectively disrupt NF-κB pathway components (e.g., RelA, IKKβ) using vectors driven by tumor-specific promoters like hTERT (48) or survivin (49). These tumor-targeted strategies offer a refined approach to safely and effectively modulate NF-κB activity in cancer therapy by concentrating drug activity within the tumor, thereby enhancing the therapeutic index and reducing systemic exposure and toxicity.

In summary, constitutive and VSV-induced NF-κB activation in PC3 prostate cancer cells drives resistance to VSV-mediated oncolysis by promoting antiviral gene expression and restricting viral replication. In contrast, VSV-sensitive LNCaP cells lack baseline NF-κB activity and antiviral defenses, allowing robust viral replication and cytolysis. Thus, NF-κB signaling is a critical determinant of resistance in aggressive prostate cancer. Targeting this pathway—while considering its molecular complexity and context-dependent roles—may sensitize resistant tumors to VSV and enhance the efficacy of oncolytic virotherapy. As such, evaluating NF-κB pathway activity in different tumor contexts may inform patient selection and guide the rational design of combination therapies aimed at overcoming viral resistance. Future studies should dissect the contributions of individual NF-κB subunits and downstream effectors and evaluate combination strategies involving NF-κB inhibition and VSV in preclinical models of advanced prostate cancer.

## Acknowledgements

We thank Dr. Kaitlin Marquis and Dr. Douglas Lyles for their support and insightful comments on data analysis and the manuscript. We thank the College of Science at the Rochester Institute of Technology (RIT) for helping to support this work. We also thank the College of Science Summer Undergraduate Research Program and the RIT Honors Program for supporting many of the undergraduates who contributed to this work. We also thank the Genomics Core at the University of Rochester for performing the RNAseq experiments. This research was supported by NIH grant 1R15CA246419 (to M. C. F.). The funders had no role in study design, data collection and interpretation, or the decision to submit the work for publication.

